# Genome sequence resources from three isolates of the apple canker pathogen *Neonectria ditissima* infecting forest trees

**DOI:** 10.1101/2024.01.13.575195

**Authors:** Salim Bourras, Heriberto Vélëz, Katarina Ihrmark, Miguel Ángel Corrales Gutiérrez, Malin Elfstrand, Larisa Garkava-Gustavsson, Kerstin Dalman Falk

## Abstract

*Neonectria ditissima* is a generalist ascomycete plant pathogen causing canker diseases on a variety of hardwood tree species and can cross-infect many of them. The fungus enters the plants through wounds throughout the year. *N. ditissima* is considered a major threat to apple production responsible for the fruit tree canker disease which damages trees and causes rotting of fruits in storage. Nearby forests and shelter belts can serve as source of inoculum for well-managed apple orchards. Thus, knowledge about the *N. ditissima* isolates infecting different host species is essential for designing integrated pest management strategies. Here, we describe the genomes of three *N. ditissima* isolates, Nd_iso34, Nd_iso35, and Nd_iso36, infecting European beech, American tulip tree, and American beech, respectively. We obtained genome assemblies of ca. 45 megabases for all isolates, covering 94% of the *N. ditissima* reference annotation, and 97% of the universal single-copy orthologs (BUSCOs). We conclude that these genome assemblies are a highly relevant resource considering the scarcity of genomic data available for *N. ditissima*.

---

The fungus, *Neonectria ditissima* (formerly *N. galligena*) causes a disease known as European canker or fruit tree canker, which is a serious threat to apple production in Sweden and globally (**Weber, 2014; Garkava-Gustavsson et al., 2016**). *N. ditissima* is a wound pathogen, able to infect trees throughout the year and fruits in orchards and storage (**Weber, 2014; Holthusen and Weber, 2021**). The fungus has a wide range of hosts (**Flack and Swinburne, 1977; Walter et al., 2015**). *N. ditissima* can attack other fruit trees such as pear (*Pyrus communis*) and quince (*Cydonia oblonga*) (**Walter et al., 2015**) as well as many different species of forest trees, e.g. *Acer, Betula, Fagus, Populus, Quercus* and *Salix* (**Ward et al., 2010**). Canker damages caused by *N. ditissima* have been recorded on forest trees in Europe including Austria, Germany, and Slovakia (**Cech, 2010; Metzler et al. 2002; Mihál, 2011**). In North America, *N. ditissima* is known to be associated with Beach Bark Disease but was also found on yellow birch (*Betula alleghaniensis*), stripped maple (*Acer pensylvanicum*) and red maple (*Acer rubrum*) (**Kasson and Livingston, 2009**). Hence, *N. ditissima* also has the propensity to cause damage to Swedish and European forestry through the extensive scarring and damage that leads to reductions in quality and value of timber (**Metzler et al., 2002**).

In this study, we sequenced three isolates, Nd_iso34, Nd_iso35, and Nd_iso36, infecting European beech (*Fagus sylvatica*), American tulip tree (*Liriodendron tulipifera*), and American beech (*Fagus grandifolia*), respectively. Fungal cultures were obtained from The Centraalbureau voor Schimmelcultures (CBS) and the Fungal Biodiversity Centre (Royal Netherlands Academy of Arts and Sciences, KNAW). Nd_iso34 (*syn*. CBS 118920, AR 3690, BPI 870951) was isolated in 2001 from European beech in Stiavnickey vrchy (Slovakia). Nd_iso35 (*syn*. CBS 118919, GJS 04-350, BPI 864075) was isolated in 2004 from American tulip tree in the Great Smokey Mountains national park (Tennessee, USA). Finally, Nd_iso36 (*syn*. CBS 118923, AR 4196, BPI 870950) was isolated in 2005 from American beech near Bass Lake (Michigan, USA).

High molecular weight genomic DNA was extracted from mycelium using the QIAGEN Genomic-tips according to the instructions from the manufacturer (QIAGEN, The Netherlands). DNA concentration and integrity were evaluated with the Qubit dsDNA Quantification Assay Kits according to the manufacturer (Invitrogen, USA), and further evaluated on 0,8 % agarose gel with Tris-acetate-Ethylenediaminetetraacetic Acid (TAE) buffer with DNA quantification standard. Sequencing libraries were prepared from 1μg DNA using the TruSeq PCRfree DNA sample preparation kit (cat# FC-121-3001/3002, Illumina Inc.) targeting an insert size of 350bp. The library preparation was performed according to the manufacturers’ instructions (guide#15036187). Libraries were then applied to 150bp paired-end sequencing on an Illumina HiSeqX sequencing system at the SNP&SEQ Technology Platform at Uppsala University (Uppsala, Sweden). The quality of the reads was checked using FASTQC, and adapters were filtered, and sequences quality trimmed using Trimmomatic v0.39 with default parameters (**Bolger et al., 2014**). Paired-end reads were then applied to ABySS, a *de novo*, parallel assembler, particularly adapted for paired-end sequences (**Simpson et al., 2009**), with default parameter. The resulting genome assemblies ranged from 44.51 Mage bases (Mb) to 45.71 Mb, with an N_50_ ranging from 200’252 to 438’025 bp (see summary in Table 1).

**Table 1.**
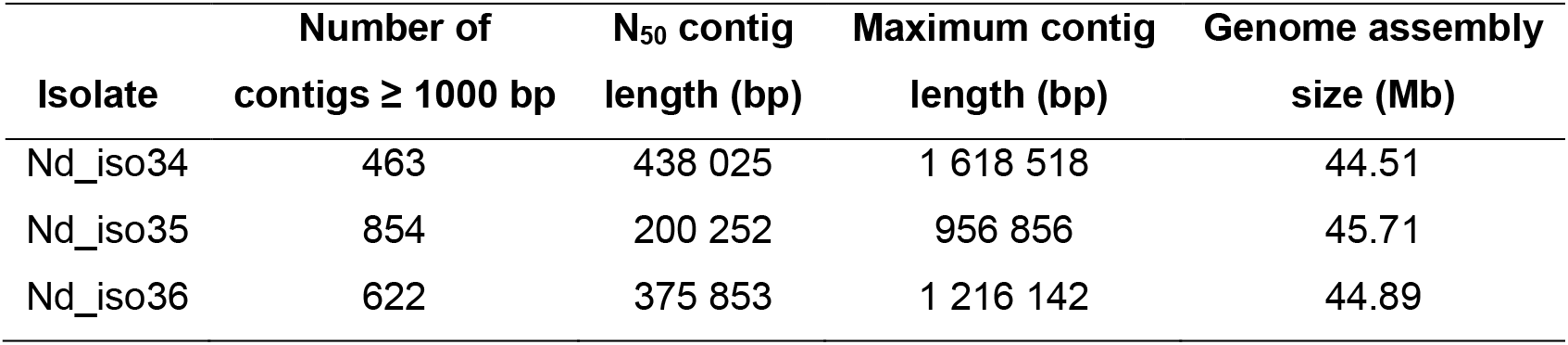
Summary of the genome assemblies of the *Neonectria ditissima* isolates Nd_iso34, Nd_iso35, and Nd_iso36.

| Isolate | Number of<br>contigs ≥ 1000 bp | N<sub>50</sub> contig<br>length (bp) | Maximum contig<br>length (bp) | Genome assembly<br>size (Mb) |
| --- | --- | --- | --- | --- |
| Nd_iso34 | 463 | 438 025 | 1 618 518 | 44.51 |
| Nd_iso35 | 854 | 200 252 | 956 856 | 45.71 |
| Nd_iso36 | 622 | 375 853 | 1 216 142 | 44.89 |

Gene prediction was carried out on contigs with a minimum size of 1000 bp (Table 1) using Funannotate v1.8.15 (**Palmer and Stajich, 2020**). A total of 11’984 (Nd_iso34), 11’890 (Nd_iso35), and 11’960 (Nd_iso36) high confidence genes were predicted, indicating that ca. 94% of the reference *N. ditissima* genome annotation (**Gómez-Cortecero et al., 2015**) is covered by our genome assemblies (Figure 1 A). Furthermore, protein size distribution was consistent over all isolates (Figure 1 B) further indicating there is no isolate-specific bias in gene completeness with respect to differences in assembly (Table 1) (**Gómez-Cortecero et al., 2015**). Finally, analysis of universal fungal single-copy orthologs (BUSCOs) (**Kim et al., 2023**) using the Funannotate-annotate pipeline, shows that ca. 97% of the BUSCO terms were recovered for isolates (Figure 1 C). We thus conclude that the resulting genome assemblies are of very good quality considering the use of short reads for genome sequencing and assembly.

**Fig. 1.**
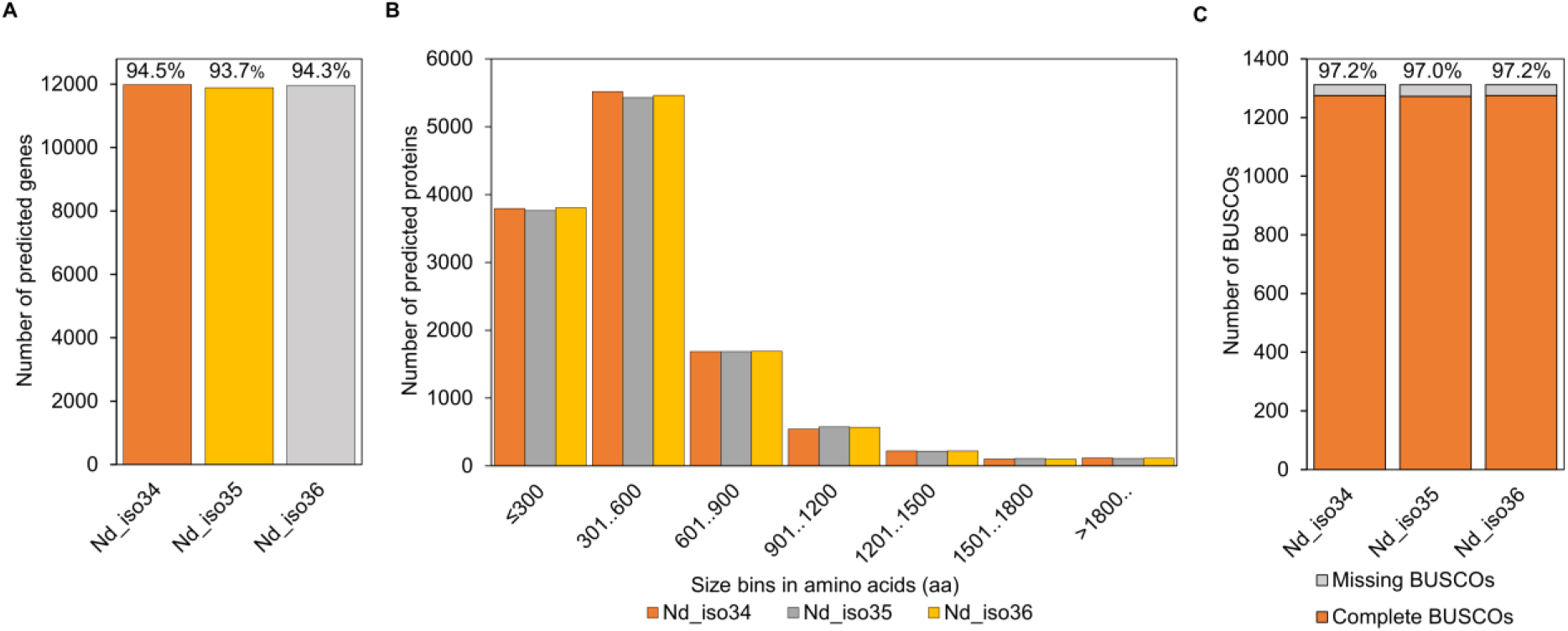
Genome annotation summary for the *Neonectria ditissima* isolates Nd_iso34, Nd_iso35, and Nd_iso36. **A**, Total number of predicted genes in every genome with the percentage of gene coverage as compared to the *N. ditissima* reference genome indicated as a percentage. **B**, protein size distribution in bins of 300 amino acids (aa). **C**, functional annotation of BUSCOs with the percentage of complete BUSCOs indicated.

Further functional annotation of the predicted proteins was performed using a combination of Funannotate-annotate v1.8.15, dbCAN (https://bcb.unl.edu/dbCAN2/), signalP v6.0 for signal peptide prediction (https://services.healthtech.dtu.dk/services/SignalP-6.0/), and targetP v.2.0 for targeting to the mitochondrion (https://services.healthtech.dtu.dk/services/TargetP-2.0/) (see summary in Table 2). Here also, results were consistent among all isolates and within the expected range with an average of 4038 protein families (Pfams), 559 Carbohydrate-Active enZymes (CAZymes), 965 putative secreted proteins with a predicted signal peptide (signalP), and 380 putative mitochondrially targeted proteins (mTP, targetP) (Table 2). We conclude that overall, the resulting genome annotations have a high level of completeness, which further demonstrates the quality and relevance of these resources for the study of *N. ditissima*.

**Table 2.** Gene functional annotation summary for the *Neonectria ditissima* isolates Nd_iso34, Nd_iso35, and Nd_iso36.

| Isolate | Number of<br>annotated<br>Pfams <sup>1</sup> | Number of<br>annotated<br>CAZymes <sup>2</sup> | Number of<br>proteins with<br>signal <sup>3</sup> | Number of<br>proteins<br>with mTP <sup>4</sup> |
| --- | --- | --- | --- | --- |
| Nd_iso34 | 4042 | 554 | 962 | 386 |
| Nd_iso35 | 4042 | 558 | 955 | 381 |
| Nd_iso36 | 4032 | 564 | 977 | 373 |

<sup>1</sup> Protein families annotated from the Pfam database.
<sup>2</sup> Carbohydrate-Active enZymes annotated from dbCAN.
<sup>3</sup> Proteins with a predicted signal peptide annotated with signalP 6.0.
<sup>4</sup> Protein with a predicted mitochondrial targeting peptide annotated with targetP 2.0.

## Data availability

Raw Illumina sequencing reads, and draft genome assemblies will be deposited at the European Nucleotide Archive (ENA, under project number PRJEB71792. Functional annotation files are made available as supplementary files as follows:

### BUSCO annotation files

- Supplementary file 1: Nd_iso34.annot.busco.txt
- Supplementary file 2: Nd_iso35.annot.busco.txt
- Supplementary file 3: Nd_iso36.annot.busco.txt

### Pfam annotation files

- Supplementary file 4: Nd_iso34.annot.pfam.txt
- Supplementary file 5: Nd_iso35.annot.pfam.txt
- Supplementary file 6: Nd_iso36.annot.pfam.txt

### dbCAN annotation files

- Supplementary file 7: Nd_iso34.annot.dbCAN.txt
- Supplementary file 8: Nd_iso35.annot.dbCAN.txt
- Supplementary file 9: Nd_iso36.annot.dbCAN.txt

## Acknowledgement

We would like to acknowledge the financial support of the Plant Protection Platform of the Swedish University of Agricultural Sciences, and the and the Carl Trygger Foundation (grant number 21:1171).

